# DNA methyltransferase inhibition induces dynamic gene expression changes in lung CD4^+^ T cells of neonatal mice with *E. coli* pneumonia

**DOI:** 10.1101/2022.09.30.510315

**Authors:** Nigel S. Michki, Roland Ndeh, Kathryn A. Helmin, Benjamin D. Singer, Sharon A. McGrath-Morrow

## Abstract

**Introduction:** Bacterial pulmonary infections are a major cause of morbidity and mortality in neonates, with less severity in older children. Previous studies demonstrated that the DNA of CD4^+^ T cells in the mouse lung, whose primary responsibility is to coordinate the immune response foreign pathogens, is differentially methylated in neonates compared with juveniles. Nevertheless, the effect of this differential DNA methylation on CD4^+^ T cell gene expression and response to infection remains unclear.

**Methods:** We treated *E. coli*-infected neonatal (4-day-old) and juvenile (13-day-old) mice with decitabine (DAC), a DNA methyltransferase inhibitor with broad-spectrum DNA demethylating activity, and performed simultaneous genome-wide DNA methylation and transcriptional profiling on lung CD4^+^ T cells.

**Results:** Juvenile and neonatal mice experienced differential demethylation in response to DAC treatment, with larger methylation differences observed in neonates. By cross-filtering differentially expressed genes between juveniles and neonates with those sites that were demethylated in neonates, we found that interferon-responsive genes such as *Ifit1* are the most down-regulated methylation-sensitive genes in neonatal mice. DAC treatment shifted neonatal lung CD4^+^ T cells toward a gene expression program similar to that of juveniles.

**Conclusion:** Following lung infection with *E. coli*, lung CD4^+^ T cells in neonatal mice exhibit epigenetic repression of important host defense pathways, which are activated by inhibition of DNA methyltransferase activity to resemble a more mature profile.

## Introduction

Pneumonia is a leading cause of death in children less than five years of age worldwide (1). The immature immune system of neonates increases the risk for adverse health outcomes. In neonates, severe pneumonias are more likely caused by bacterial rather than viral pathogens, since transfer of maternal antibodies against viral pathogens confers some protection to the neonate (2–5). Treatments for neonatal pneumonia are currently limited to antibiotics and supportive care, and mortality remains unacceptably high.

In older children, maturation of the adaptive immune system occurs due to both thymic development and the accumulation of antigenic experiences. Consequently, older children have more efficient and robust immune responses when challenged with bacterial pathogens and lower morbidity and mortality compared with neonates. Modulation of the neonatal immune response toward a more mature effector T cell profile could improve early immune responses to bacterial infections in the neonate, yet interventions that accelerate maturation of the neonatal immune system are not available.

We previously reported differential expression of immune effector genes in neonatal lung CD4^+^ T cells compared with juveniles in response to *E. coli* pneumonia. Furthermore, we found that the loci encoding immune effector genes in neonates were more likely to exhibit persistent DNA methylation in response to *E. coli*. Using a computational approach, we found that demethylation of immune effector genes in response to *E. coli* challenge correlated with increased gene expression (4). We also observed that treatment of mice with a DNA methyltransferase inhibitor resulted in changes to lung CD4^+^ T cell phenotype that were more robust in neonates than juveniles infected with *E. coli*.

In this study, we sought to determine whether administration of a DNA methyltransferase inhibitor with broad-spectrum hypomethylating activity, decitabine (DAC), to neonates post-*E. coli* challenge would induce hypomethylation of immune effector genes in lung CD4^+^ T cells to shift their gene expression profile toward a more mature (juvenile) state. We administered DAC to neonates and juveniles following intrapharyngeal aspiration of *E. coli*. Using a computational approach, we found that immune effector genes, including core genes involved in interferon signaling, in lung CD4^+^ T cells sorted from DAC-treated neonates infected with *E. coli* underwent significant demethylation that was associated with gene expression changes. The resulting gene expression signatures resembled those of juvenile lung CD4^+^ T cells post-*E. coli*. This study indicates that in a model of neonatal pneumonia, DNA methyltransferase inhibition modulates dynamic gene expression in lung CD4^+^ T cells toward a more mature T cell phenotype.

## Methods

### Mice

Timed pregnant C57BL/6NJ mice were obtained from Charles River Laboratories. Adult animals were maintained on an AIN 76A diet, given water *ad libitum*, and housed at a temperature range of 20-23°C under 12-hour light/dark cycles. Pups of both sexes were used in the reported experiments. Experiments were conducted in accordance with the standards established by the United States Animal Welfare Act set forth in National Institutes of Health guidelines and the Policy and Procedures Manual of the Johns Hopkins University Animal Care and Use Committee.

### Intrapharyngeal aspiration of *E. coli*

Pups were sedated with isoflurane prior to intrapharyngeal aspiration with *E. coli* bacteria (Seattle 1946, serotype O6, ATCC 25922). Neonatal (4-day-old) and juvenile (13-day-old) mice were randomized by cage to receive either PBS alone or *E. coli* in PBS (2.4 × 10^6^ colony-forming units). Bacterial aspiration and subsequent visualization was performed as described previously (6). Neonatal mice were aspirated with 10 μL of fluid and juvenile mice with 15 μL of fluid.

### Quantitative microbiology of *E. coli* bacteria

*E. coli* bacteria were streaked on an LB agar plate and grown overnight at 37°C. Bacteria were transferred to LB medium, agitated at 250 RPM, and incubated at 37°C for 3-4 hours. Bacterial growth was determined using optical density (OD) measured at 600 nm, with serial dilutions performed and plated overnight at 37 °C to assess OD measurement accuracy.

### Decitabine administration

Neonatal and juvenile mice received daily intraperitoneal injections of 5-aza-2’-deoxycytidine (decitabine, DAC; Sigma) 1 mg/kg (7) in 30 μL of diluted DMSO or diluted DMSO alone, beginning 24 hours and 48 hours following aspiration of 2.4 × 10^6^ colony-forming units of *E. coli* in PBS. Mice were euthanized 72 hours post-*E. coli* aspiration.

### Processing of mouse lungs

Preparation of lung tissue for histology and processing to create single-cell suspensions were performed as previously reported (7–9). Briefly, lung tissue was minced in a petri dish with a buffer containing DNase and collagenase I and incubated at 37°C for 30 minutes. The resulting tissue suspension was mechanically disrupted by passing through an 18-gauge needle and filtered through a 70-μm pore size cell strainer. PBS was added before centrifugation at 300 x g for 10 minutes. ACK lysing buffer was added to the pellet, incubated at room temperature for 5 minutes, and quenched using PBS. The suspension was finally filtered, spun at 300xg for 5 minutes, and resuspended in MACS buffer (PBS + 0.5% bovine serum albumin + 2 mM EDTA). Neonatal samples were each prepared using lungs from three separate mice subjected to the same conditions from 3-4 different litters. Juvenile samples were prepared using lungs from two separate mice subjected to the same conditions from 2-3 different litters.

### CD4^+^ T-cell positive selection from mouse lungs

CD4^+^ T-cell isolation was performed as previously described (10). Briefly, lung cell suspensions were incubated with CD4-PE conjugated antibody (BD Biosciences) at 4°C for 10 minutes before washing with MACS buffer, pelleting at 300xg for 10 minutes, and resuspending in MACS buffer + 20 μL of anti-PE microbeads (Miltenyi Biotec). Cells were then magnetically separated using an MS column (Miltenyi Biotec) per the manufacturer’s instructions, suspended in a sorting medium (PBS + 0.5% BSA, 0.5% fetal bovine serum, 1 mM EDTA, and 25 mM HEPES) and placed on ice before sorting. Lung CD4^+^ T-cells were sorted based on characteristic low FSC and SSC and PE^+^ status using a BD FACSAria II instrument with FACSDiva software (BD). Sorted cells were pelleted at 5000 x g for 10 minutes, the supernatant was removed, and the cell pellet was lysed with 350 μL of buffer RLTplus (Qiagen) supplemented with 1% β-mercaptoethanol for 5 minutes before storage at -80 °C.

### Modified reduced representation bisulfite sequencing

Genomic DNA isolation and mRRBS was performed as previously described (4, 11, 12). Briefly, gDNA was isolated using the AllPrep DNA/RNA Micro Kit (Qiagen) and quantified with a Qubit 3.0 instrument. Approximately 50-200 ng of gDNA was digested with the restriction endonuclease MspI (New England BioLabs) per the manufacturer’s recommendations. Resulting fragmented DNA underwent size selection for fragments approximately 100-250 bp in length using SPRI beads (MagBio Genomics) and were subsequently subjected to bisulfite conversion using the EZ DNA Methylation-Lightning Kit (Zymo Research). Libraries for Illumina-based sequencing were prepared with the Pico Methyl-Seq Library Prep Kit (Zymo Research) using Illumina TruSeq DNA methylation indices. Libraries were run on a High Sensitivity chip using an Agilent TapeStation 4200 to assess size distribution and overall quality of the amplified libraries. Pooled libraries were sequenced on an Illumina NextSeq 500 instrument using the NextSeq 500/550 V2 High Output reagent kit (1 × 75 cycles).

Indexed samples were demultiplexed to fastq files with BCL2FASTQ v2.17.1.14. Reads were trimmed of 10 base pairs from the 5’ end with TrimGalore! v0.4.3. Sequence alignment to the GRCm38/mm10 reference genome and methylation extraction ignoring 1 base at the 3’ end (after reviewing the M-bias plots) were performed with Bismark v0.16.3. Bismark coverage (counts) files for cytosines in CpG context were analyzed with respect to differential methylation with the protocol outlined in (13). Briefly, methylation frequency (how often a CpG position is methylated, termed β) was calculated from the coverage files generated by Bismark using the read.bismark function from the R package DSS (14), with *loci=NULL, rmZeroCov=FALSE, strandCollapse=FALSE*, and *replace=TRUE*. Tests for differential methylation at each CpG position were performed within age groups and between DAC and DMSO treatments using the DMLtest function, with *smoothing=FALSE* and *equal*.*disp=FALSE*. CpG positions showing differential methylation were filtered to only include those with FDR < 0.05. Differentially methylated CpG positions were annotated with extra genomic information using HOMER v4.11 (15), in particular to determine the distance of each CpG to the nearest gene’s transcriptional start site (TSS).

### RNA-sequencing

Nucleic acid isolation and library preparation were performed as previously described (4), using the QIAGEN AllPrep DNA/RNA Micro Kit and the SMARTer Stranded Total RNA-Seq Kit, version 2 (Takara). Fastq files were generated from bcl files using BCL2FASTQ v2.17.1.14 with default parameters. Reads were aligned to the NCBI mouse genome (mm10/GRCm38) using the nf-core/rnaseq v3.8.1 Nextflow pipeline (16, 17). Briefly, in this pipeline, reads were trimmed using TrimGalore!, a wrapper around Cutadapt (18) and FastQC, was used to remove sequencing adapters. Trimmed reads were aligned to the genome using STAR 2.7.10a (19) and quantified at the gene level using Salmon 1.9.0 (20) to generate a gene-by-sample counts matrix. Differential gene expression analysis was performed using DESeq2 (21). Differentially expressed genes were identified between groups using the standard DESeq2 results test with no additional shrinkage estimators applied and the model design formula *f ∼ age + treatment + age:treatment*. DEGs were called with a false-discovery rate (FDR) cutoff of FDR < 0.05 and a log_2_-fold change (LFC) cutoff of magnitude

1. Variance stabilized counts used for plotting and tertiary analysis were generated using the *rlog* function with *blind=FALSE*.

### Statistical analysis

Identification of genes simultaneously differentially methylated at the DNA level and differentially expressed at the RNA level was accomplished using a cross-filtering strategy. Beginning with all genes annotated in the mouse genome, genes were first filtered by determining if, in the neonatal cohort, at least one CpG located either intragenically or within 2.5 kb up/downstream of the TSS of the gene was differentially methylated with DAC treatment. These genes were then further filtered based on their differential gene expression as described previously (4). In order to rank these cross-filtered differentially methylated and differentially expressed genes the *rank_genes_groups* function from scanpy (22) was used on the previously rlog-transformed gene expression values with *method=‘wilcoxon’*. The top differentially expressed genes identified here with an FDR < 0.05 were analyzed using the Statistical Overrepresentation Test from PANTHER v17.0 (23). Genes ordered by FDR were analyzed against GO: Biological Processes using all annotated *mus musculus* genes as background, with *test type = Fisher’s Exact* and *Correction = Calculated False Discovery Rate*. The top genes (above) were also analyzed using the gProfiler (24) functional profiling tool in order to identify transcription factor binding motifs that were enriched in their promoter regions. Default runtime options were used (*unordered query, only annotated genes, g:SCS threshold < 0*.*05*).

### Data availability

The raw and processed next-generation sequencing data sets have been uploaded to the GEO database (https://www.ncbi.nlm.nih.gov/geo/) under accession number GSE214490, which will be made public upon peer-reviewed publication.

## Results

### DAC administration induces dynamic alterations in epigenetic and transcriptional signatures within neonatal and juvenile lung CD4^+^ T cells

Neonatal and juvenile mice were challenged with *E. coli* by intrapharyngeal aspiration. Following *E. coli* aspiration, neonatal and juvenile mice received intraperitoneal injections at 24 and 48 hours with either decitabine (DAC) or DMSO (control). Lung CD4^+^ T cells were then harvested at 72 hours post-*E. coli* aspiration and bulk mRRBS and RNA-seq were performed on isolated DNA and RNA, respectively (**Figure 1A**).

**Figure 1:**
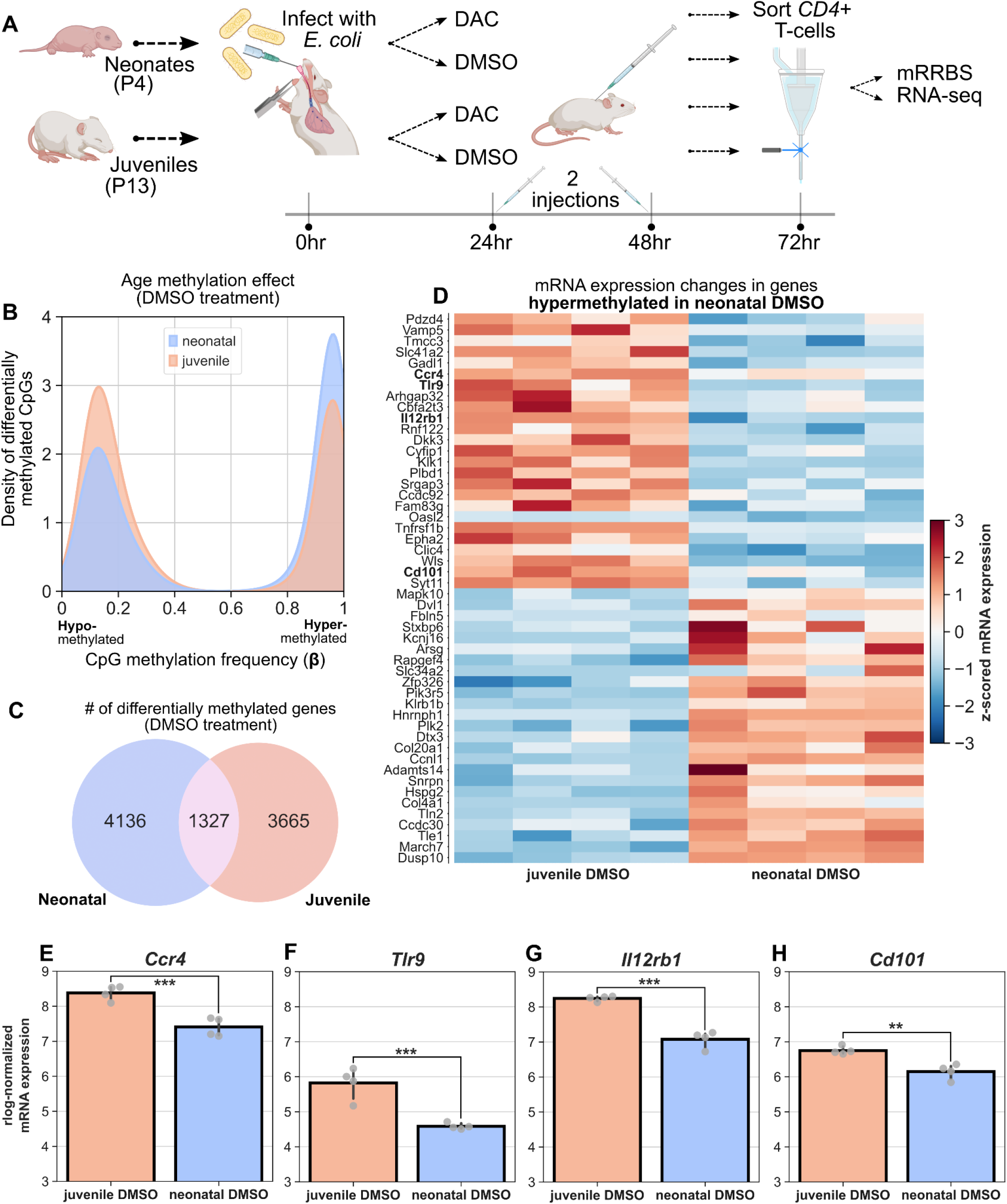
Age-related changes in CpG methylation and mRNA expression in *E. coli* infected mice. **(A)** Experimental overview. Neonatal (P4) and juvenile (P13) mice were infected with *E. coli* via intrapharyngeal aspiration before treatment with either DAC or DMSO at 24 and 48 hours post-infection. At 72 hours post-infection, mice were sacrificed and lung CD4^+^ T-cells were sorted for bulk mRRBS and RNA-seq. **(B)** Density plot of methylation level (β) across all differentially methylated CpGs in *E. coli*-infected juvenile and neonatal DMSO-treated mice at 72 hours post-infection. CpGs in neonates are more likely to be methylated than those in juveniles. **(C)** Venn diagram showing the number and overlap of differentially methylated genes with at least 1 differentially methylated CpG in neonates versus juveniles. **(D)** Matrixplot of top genes over-/under-expressed in juveniles versus neonates that are also hypermethylated in neonates. Expression levels are z-scored. Bolded genes are related to immune processes. Figure restricted to top 25 genes for visibility. **(E-H)** *rlog*-normalized mRNA expression levels of *Ccr4, Tlr9, Il12rb1*, and *Cd101*, four immune-process related genes. Two asterisks indicate FDR < 0.005, three asterisks indicate FDR < 0.0005.

We began by comparing CpG methylation profiles in juvenile versus neonatal mice infected with *E. coli* and treated with DMSO. After processing and quality filtering, we found that 12,779 CpGs were hypermethylated in neonatal versus juvenile mice and 9,240 CpGs were hypermethylated in juveniles versus neonates. The distribution of methylation frequencies (β) across all differentially methylated CpGs was skewed toward hypermethylation in neonates (**Figure 1B**). 4,136 genes in neonatal mice had at least one hypermethylated CpG that was either intragenic or within 2.5 kb of its TSS. 3,665 genes were hypermethylated in juveniles and 1,327 genes were shared between age groups (**Figure 1C**). This hypermethylation of neonatal lung CD4^+^ T cells relative to juveniles recapitulates our previous study in *E. coli*-infected mice at 48 hours post-infection (4).

We next evaluated mRNA expression changes between juvenile and neonatal *E. coli*-infected mice treated with DMSO. Restricting our analysis to include only those genes with hypermethylated CpGs in neonates that were intragenic or within 2.5 kb of their TSS (**Figure 1D**), we ranked the top differentially genes between the two DMSO-treated age groups out of the 99 genes that were simultaneously differentially expressed between juveniles and neonates and were hypermethylated in neonates. These genes were associated with diverse biological processes, however *Ccr4, Tlr9, Il12rb1*, and *Cd101* were all simultaneously up-regulated in juveniles, hypermethylated in neonates, and associated with important immune processes (**Figure 1E-H**), indicating that in an infected, DMSO-treated state, the expression of these genes may be directly repressed by age-related epigenetic marks.

Examination of the response of *E. coli*-infected mice to DAC treatment at the DNA methylation level revealed that the frequencies of hypo- and hyper-methylation were markedly different between age groups in response to DAC administration. Neonatal lung CD4^+^ T cells were more responsive to DAC, with DAC-treated neonatal CD4^+^ T cells exhibiting a greater frequency of hypomethylation than DAC-treated juveniles (**Figure 2A**,**B**), which had similar frequencies of CpG hypo- and hyper-methylation. These findings indicate that neonates experience a greater degree of demethylation than juveniles in response to DAC administration.

**Figure 2:**
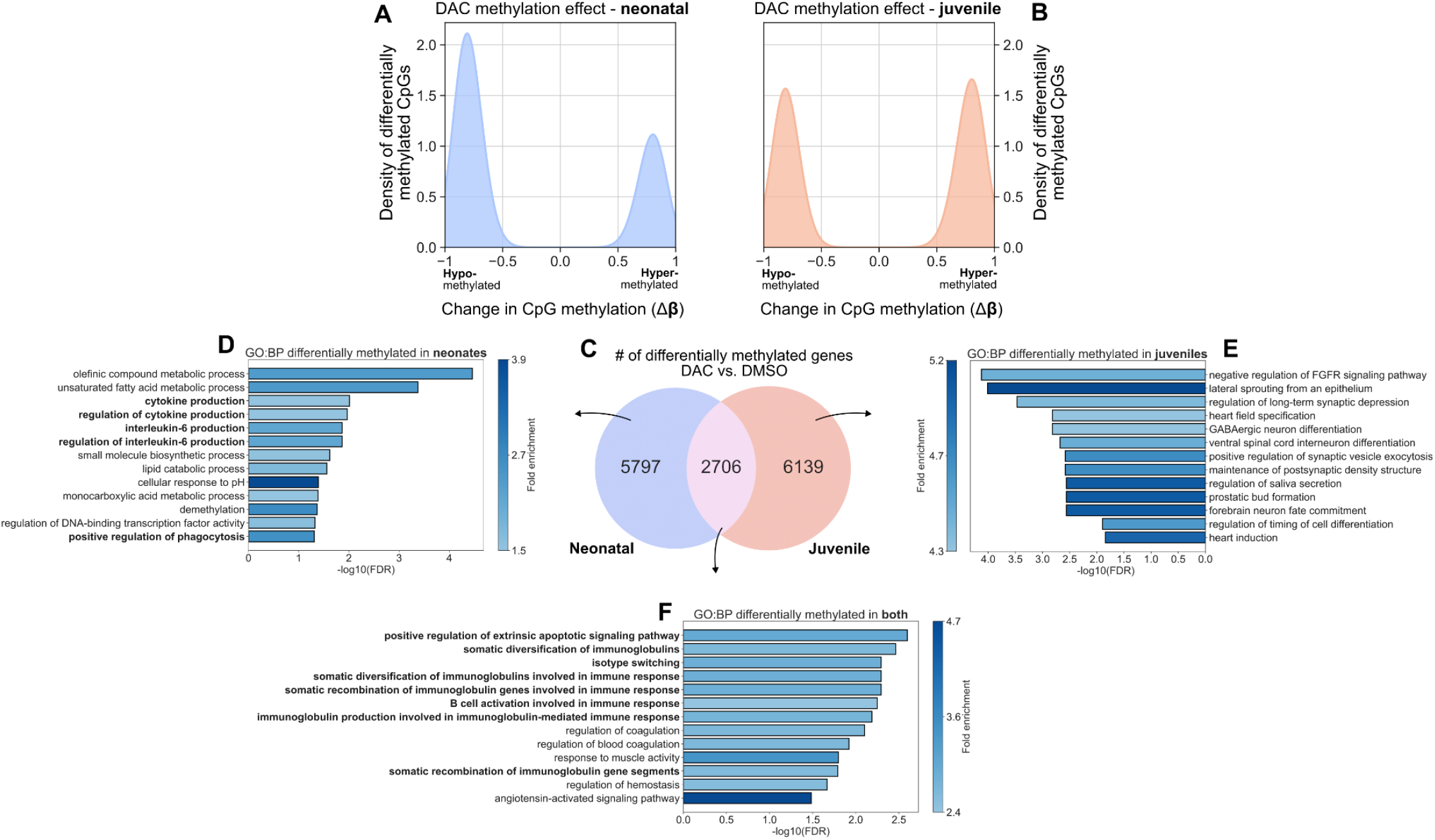
DAC treatment of mice infected with *E. coli* induces CpG demethylation at unique genomic loci and with greater strength in neonates versus juveniles. **(A**,**B)** Density plot of changes in CpG methylation frequency (Δβ) in (**A**) neonatal mice and (**B**) juvenile mice. DAC treatment more strongly induces demethylation (Δβ < 0) in neonates than in juveniles, which are less likely to be demethylated (Δβ > 0). **(C)** Venn diagram showing the number and overlap of differentially methylated genes with at least one differentially methylated CpG in DAC vs. DMSO across neonates and juveniles. **(D-F)** Over-representation analysis (ORA) from gProfiler showing GO: Biological Processes (GO:BPs) associated with genes differentially methylated in response to DAC treatment in (**D**) neonates, (**E**) juveniles, and (**F**) both.

We then asked which genes were differentially methylated with DAC administration. We considered a CpG to be associated with a specific gene if it was contained within the annotated gene locus or within 2.5 kb of its TSS, and we considered a gene to be differentially methylated if at least one CpG associated with it was differentially methylated. We further remained naïve to the direction of differential methylation (hypo-versus hyper-methylated) in response to DAC administration, as the link between CpG hypo- and hyper-methylation and gene expression is not always obvious (13, 25, 26). We identified 5,795 genes that were differentially methylated in neonates, 6,739 in juveniles, and 2,706 in both age groups in response to DAC administration (**Figure 2C, Supplemental Table 1**). In neonatal mice, over-represented GO: Biological Processes (GO:BPs) included *Unsaturated fatty acid metabolic process, Positive regulation of transport*, and *IL-6 production* (**Figure 2D**), whereas in juveniles, over-represented GO:BPs included *Cell morphogenesis involved in differentiation* and *Cell-cell communication* (**Figure 2E, Supplemental Tables 2-4)**. Over-representation analysis (ORA) of GO:BPs in genes differentially methylated in both age groups included *Positive regulation of signal transduction, Cellular response to stress*, and *Response to extracellular stimulus* (**Figure 2F**). Collectively, these results indicate that DAC administration results in altered DNA methylation and transcriptional signatures within both neonatal and juvenile lung CD4+ T cells, with a greater demethylation effect in neonates.

### DAC induces dynamic expression of genes associated with immune responses in neonatal lung CD4^+^ T cells post-*E. coli*

Transcription is the proximate readout of epigenetic control mechanisms (27). Accordingly, we asked whether there were gene expression changes in response to DAC treatment and whether these changes differed with age. Using DESeq2 (21), we independently regressed out the contribution of age, treatment, and the interaction between the two in order to identify which genes demonstrated the most marked expression changes (|LFC| > 1) with respect to each (**Figure 3A-C, Supplemental Tables 5-7**). We likewise determined which specific biological processes were associated with these gene sets, and using gProfiler (24) performed an ORA that revealed genes involved in *host defense responses* were up-regulated as a result of DAC administration and that neonates experienced larger expression changes in these gene programs than juveniles (**Figure 3D-I**).

**Figure 3.**
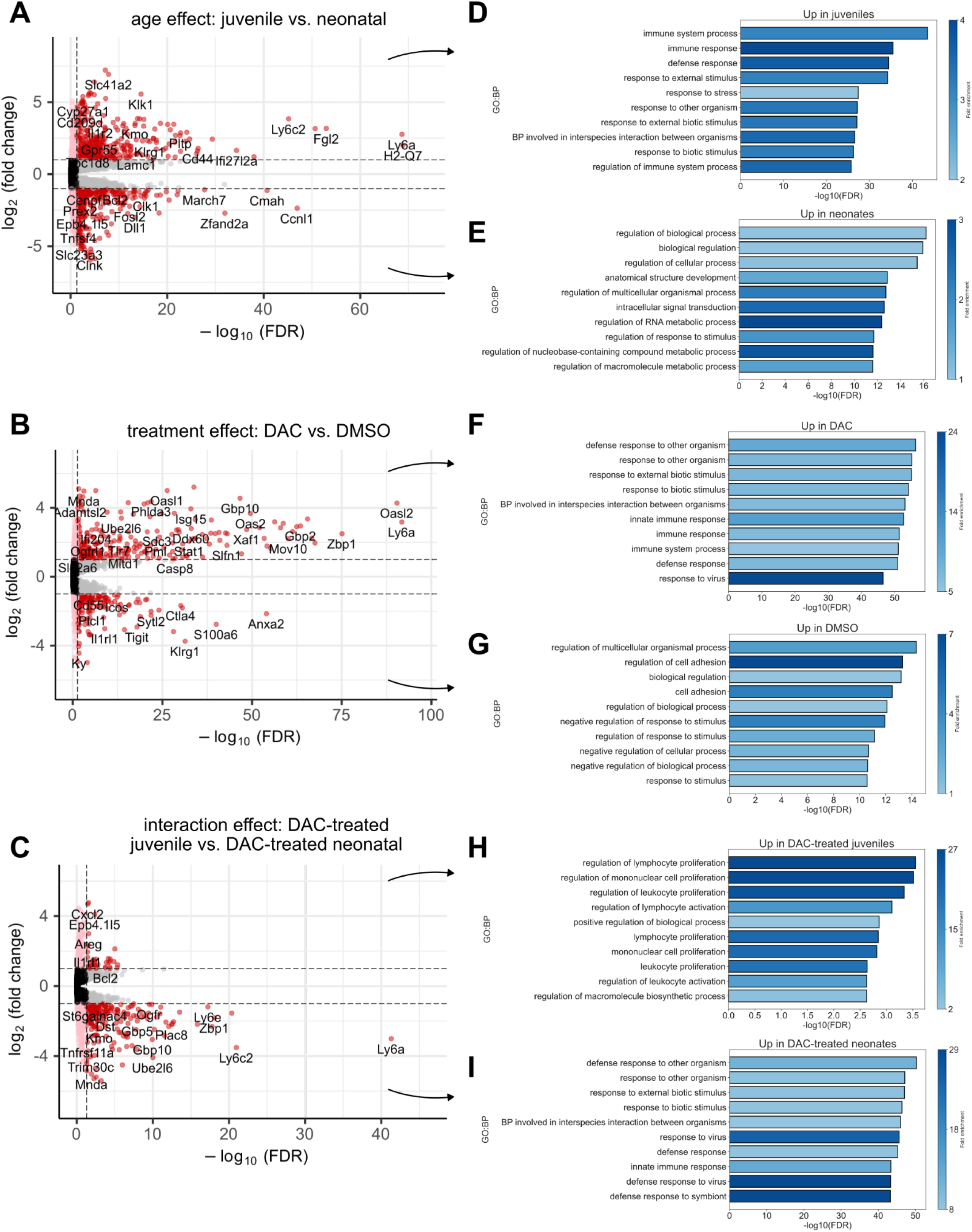
Age and DAC treatment independently and in concert induce gene expression changes in immune program genes. **(A)** Volcano plot showing the log_2_-fold changes (LFCs) in gene expression as a function of age (juvenile versus neonatal). Genes with positive LFC values are expressed at higher levels in juveniles than neonates. **(B)** As in (**A**) for the DAC versus DMSO comparison after regressing out the effect of age. **(C)** As in (**A**,**B**) for the interaction between age and treatment status. **(D)** Over-representation analysis (ORA) from gProfiler showing GO: Biological Processes (GO:BP) associated with genes up-regulated in juveniles versus neonates, and **(E)** neonates versus juveniles. Fold-enrichment (colorbar) in this context is the number of genes associated with a term divided by the number of genes expected to be associated with that term by random chance. **(F)** as in (**D**) for genes up-regulated CD4^+^ T cells from DAC-treated mice versus DMSO, and **(G)** for DMSO-treated mice versus DAC. **(H)** as in (**D**) for genes more greatly increased in expression in DAC-treated juveniles than neonates. **(I)** as in (**E**) for genes more greatly increased in expression in DAC-treated neonates than juveniles.

We first considered how age and DAC administration independently contributed to gene expression changes observed in our experiment. Genes associated with immune system and host defense processes are expressed at significantly higher transcript levels in juveniles versus neonates, consistent with previous reports (28) (**Figure 3A**,**D**). Irrespective of age, administration of DAC similarly increased the expression of genes associated with immune responses, with marked increases in expression of *Zbp1, Ifit3, Igtp*, and *Isg15*, indicating that interferon response programs are particularly affected by DAC administration (**Figure 3B**,**F**). Interestingly, genes associated with cell adhesion appeared to be down-regulated in response to DAC administration, suggesting a cell migratory phenotype. Likewise, genes associated with the *negative* regulation of response to stimuli were down-regulated in response to DAC administration, suggesting that cells exposed to DAC may be *more* responsive to external stimuli (**Figure 3B**,**G**).

We next considered the interaction between age and DAC administration on gene expression changes. Genes with positive LFCs with respect to this interaction term were those demonstrating large increases in expression in DAC-treated juveniles but less in DAC-treated neonates (**Figure 3C**,**I**), and those with negative LFCs exhibited large increases in expression in DAC-treated neonates but less in DAC-treated juveniles (**Figure 3C**,**H**). Few genes were up-regulated to a greater degree in DAC-treated juveniles than neonates and were not a primary focus of our analysis, though those that were up-regulated were involved in the regulation of *Lymphocyte development* and *Proliferation*. In contrast, biological processes up-regulated more in DAC-treated neonates included *Defense response to other organism, Response to external biotic stimulus*, and *Innate immune system response*. We call particular attention to *Zbp1, Tap1*, and *Oas1a*, three genes that are directly involved in interferon response pathways. *Zbp1* encodes a Z-DNA binding protein that classically binds to foreign DNA to induce type-I interferon production (29). It additionally mediates interferon-induced necroptosis via its interaction with RIPK3 (30). *Tap1* encodes a membrane-bound antigen transporter responsible for transporting peptides across the endoplasmic reticulum for MHC-I molecule assembly (31, 32). Itself an interferon-stimulated gene, *Tap1* can additionally stimulate the production of IFN-β (33). *Oas1a* encodes 2’-5’-oligoadenylate synthetase 1, an interferon-inducible protein that, upon binding to dsRNA, generates 2’-5’ oligoadenylates that activate *RNaseL (34)*, an antiviral endoribonuclease which is also essential for antibacterial immunity (35), (36). Taken together, these data indicate that expected age-related changes in gene expression related to immunity are recapitulated in CD4^+^ T cells from our experiment, that DAC administration appears to increase the expression of genes related to interferon response programs, and that neonates are particularly sensitive to DAC-induced increases in expression of genes that are critical modulators of immune defense responses to pathogens.

### DAC induces juvenile-like expression of interferon program genes in neonatal lung CD4^+^ T cells

To determine which differentially methylated genes were also differentially expressed in response to DAC administration, we filtered genes detected in our bulk RNA-seq samples to include only those that were differentially methylated in response to DAC treatment in neonatal mice. Subsequently, we performed differential expression tests on the mRNA expression levels from neonatal DMSO-treated mice with those from all other cohorts combined (juvenile DAC, juvenile DMSO, and neonatal DAC) to identify those genes whose expression levels shifted to an expression level similar to that of DMSO- or DAC-treated juvenile mice (**Figure 4A**,**B**). Genes up-regulated in neonatal mice following DAC administration included *Zbp1, Ifit1, Psmb10, Igtp*, and *Isg20* (**Figure 4B, bold**), which are involved in interferon signaling. *Zbp1* encodes a Z-DNA binding protein that binds to foreign DNA to induce type-I interferon production (29) and mediates interferon-induced necroptosis via its interaction with RIPK3 (30). *Ifit1* (*Isg56*) is an interferon-stimulated gene whose protein product exerts a modulatory role on the interferon gene program, shifting LPS-exposed macrophages away from an interferon gene program and toward a non-inflammatory one, thereby limiting their bacterial susceptibility (37). *Psmb10* codes for a component (*MECL1/ 2i*)of the immunoproteasome, a protein complex involved in antigen processing and presentation, and expression of *Psmb10* is known to be stimulated by interferon-γ (38). *Igtp* (*Irgm3*) is an interferon-γ-stimulated GTPase that affects host defense in a complex, pathogen-dependent manner (39) in concert with other interferon-stimulated genes such as *Irgm1* (40). Finally, *Isg20* encodes an interferon-stimulated 3’-5’ ribonuclease which has been extensively studied in the context of RNA-viruses but may also play a role in innate antibacterial defense responses (41–43).

**Figure 4.**
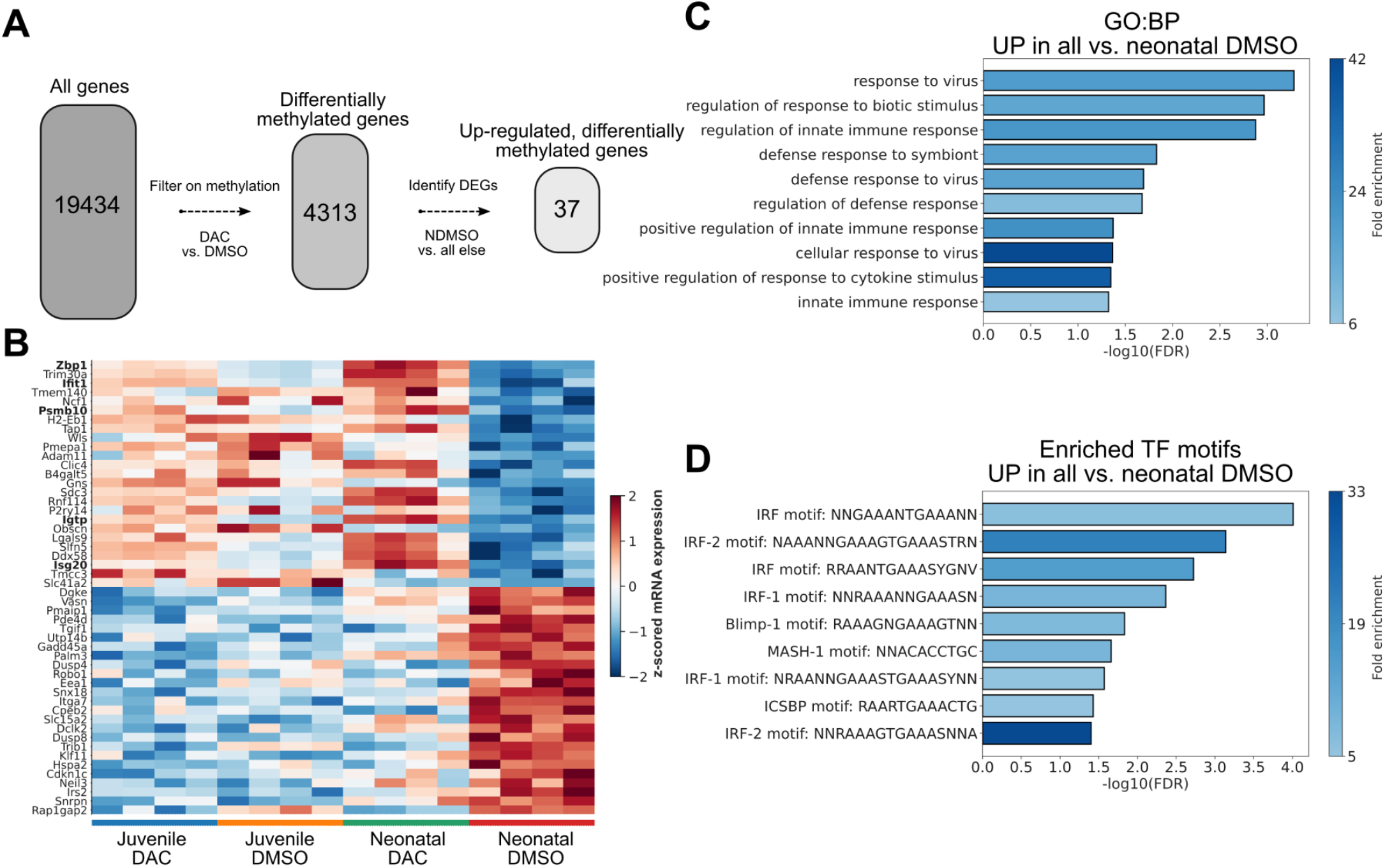
DAC treatment of neonatal mice infected with *E. coli* induces a juvenile-like immune response in CD4^+^ T cells. **(A)** Multi-level computational filtering overview. Genes were filtered to include only those that show a change in methylation state in response to DAC treatment in either age group. These differentially methylated genes were then ranked by their differential mRNA expression when comparing neonatal DMSO to all other groups. **(B)** Matrixplot of scaled mRNA expression levels for the top DEGs in each direction in the comparison of neonatal DMSO versus all else. Bolded genes are directly involved in type-I interferon responses. Figure restricted to top 25 genes for visibility. **(C)** Over-Enrichment Analysis (ORA) of GO:BPs terms for the top DEGs identified in (**B**), showing over-representation of terms related to viral and bacterial defense responses. **(D)** ORA as in (**C**) for transcription factor (TF) motifs, showing over-representation of interferon response factor (IRF) motifs in the promoters of up-regulated DEGs.

We then determined whether any specific biological processes were coherently associated with genes up-regulated in response to DAC administration in neonatal mice. Using an ORA from PantherDB (23) to characterize the up-regulated genes, we identified a number of GO:BPs associated with these genes (**Figure 4C**), with top terms including *Response to virus, Regulation of response to biotic stimulus*, and *Regulation of innate immune response*. Furthermore, we asked whether any specific transcription factor (TF) motifs were over-represented in the promoter regions of the up-regulated genes, and using gProfiler (24) identified *IRF*-motifs as being particularly over-represented (**Figure 4D**). We performed similar analyses for genes down-regulated in response to DAC administration and found that the GO:BP terms *Negative regulation of cell-cell signaling, negative regulation of response to stimulus*, and *Phosphorus metabolic process* were all over-represented in that gene set. Likewise, the TF-motifs KLF3, GKLF, and BTEB2 were over-represented in the promoter regions of genes down-regulated in response to DAC. The transcription factor KLF3 is interesting in this context, as it suppresses NF-κB–driven inflammation in mice (44), indicating that its down-regulation in response to DAC may support a pro-inflammatory anti-bacterial phenotype in concert with some of the interferon-stimulated genes discussed previously (e.g., *Ifit1*). Taken together, these data indicate that lung CD4^+^ cells in neonatal mice treated with DAC are differentially methylated at the loci of genes known to be involved in the balance between interferon and inflammatory responses, and these genes shift in expression at the transcript level to promote a pro-inflammatory phenotype more similar to that of juvenile mice.

## Discussion

In this study, inhibition of DNA methyltransferase activity with DAC induced dynamic gene expression in neonatal lung CD4^+^ T cells in a model of *E. coli* pneumonia to a greater extent than that found in older (juvenile) mice. These gene expression changes post-DAC treatment highlight the role of DNA methylation in regulating immune response genes in an age-specific manner. Integrating methylation and bulk RNA-seq expression data using a multi-modal informatics approach, we determined that interferon pathway genes were particularly sensitive to DAC-induced demethylation in the neonate. In contrast, DAC-treated juvenile CD4^+^ T cells in *E. coli*-infected lungs demonstrated only modest gene expression changes compared with neonates. Neonatal susceptibility to DNA methyltransferase inhibition enabled neonatal lung CD4^+^ T cells to exhibit transcriptional plasticity toward a juvenile-like response to *E. coli*.

We previously reported that neonates challenged with *E. coli* do not exhibit hypomethylation of key immune response genes to the same extent as juveniles (4). Whereas neonates challenged with *E. coli* had attenuated hypomethylation of CpGs, juveniles were poised to quickly respond to *E. coli* with dynamic hypomethylation of CpGs and marked induction of *Stat1* and interferon response genes. When neonates were treated with DAC following *E. coli* challenge, however, they exhibited marked hypomethylation of several immune response genes, including those associated with interferon pathways and an effector T cell phenotype. The results presented here implicating epigenetic repression of interferon pathway genes echo epigenetic mechanisms discovered in zebrafish embryos, where conservation of DNA methylation occurs through repression of transposons that prevent activation of interferon response genes (45). In the zebrafish model, the loss of DNA methylation through mutations in *Uhrf1* or *Dnmt1* occurred in genes critical to the interferon pathways. Similarly, in our system, DAC led to hypomethylation of CpGs associated with interferon pathway genes in neonatal lung CD4^+^ T cells.

That CpGs in neonatal lung CD4^+^ T cells were more frequently hypomethylated than juveniles in response to DAC administration implies that juvenile CD4^+^ T cells have more robust epigenetic maintenance mechanisms in place to stabilize CpG methylation in response to a demethylating agent such as DAC. Methylation maintenance mechanisms are not fully understood, but previous evidence suggests that there is a mutual relationship between CpG methylation and the recruitment of DNA methyltransferase proteins such as DNMT3A/B (46–49). Mammalian genomes become progressively hyper-methylated over the life course (50, 51), suggesting that neonatal mice may lack effective methylation maintenance machinery, which results in phenotypic plasticity.

There are limitations to our model. The clinical utility of DAC administration in neonatal pneumonia is unclear and was not the intent of our study. DAC may have off-site effects caused by its systemic administration. Future studies examining the feasibility of aerosolized DAC for local administration into the lungs may be warranted. DAC has also been shown to inhibit cell growth of lung CD4^+^ T cells in neonate and juvenile mice (4), which may negate the benefits of inducing a CD4^+^ T cell effector phenotype and interferon pathways needed to combat bacterial pathogens, such as *E. coli*. Nevertheless, our study suggests that DNA methyltransferase inhibition induces CD4^+^ T cell plasticity in the lung cells of neonates to activate transcriptional programming resembling the mature response to infection.

## Supplemental Tables

**Supplemental Table 1**: Genes differentially methylated in response to DAC treatment across ages.

**Supplemental Tables 2-4:** gProfiler ORA analysis results for genes differentially methylated in (**2**) juveniles, (**3**) neonates, and (**4**) both.

**Supplemental Tables 5-7**: DEseq2 differential expression test results for the (**5**) effect of age (juvenile versus neonatal), (**6**) effect of treatment (DAC versus DMSO), and (**7**) marginal interaction effect between age and treatment.

